# Impact of autologous serum on an *in vitro* granuloma model to study *Mycobacterium tuberculosis* infection dynamics

**DOI:** 10.1101/2024.12.27.630364

**Authors:** Chloé Bourg, Elisabeth Hodille, Florence Ader, Oana Dumitrescu, Charlotte Genestet, the Lyon TB study group

## Abstract

Tuberculosis (TB) remains a global health challenge, with *Mycobacterium tuberculosis* (Mtb) persisting in granulomas, complex immune structures that balance containment and survival of the pathogen. *In vitro* granuloma models have advanced TB research but often rely on commercial human serum, which may not fully replicate the native immune environment. Here, we investigated the role of autologous serum, either heat-inactivated or native, in granuloma-like structure (GLS) formation, bacterial dynamics, and immune responses. Using a human PBMC-based model to induce spontaneous *in vitro* granuloma formation after infection with Mtb, we found that autologous serum significantly enhanced GLS formation and altered cytokine profiles compared to commercial serum. Native serum also reduced bacterial growth, likely due to active complement system-mediated immune responses. These findings highlight the importance of autologous serum in creating physiologically relevant models, crucial for preclinical evaluation of TB therapies. Incorporating patient-specific serum can improve model accuracy, enabling better exploration of host-pathogen interactions and targeted therapeutic strategies.

## Dear Editor

Tuberculosis (TB) remains a significant global health challenge, with *Mycobacterium tuberculosis* (Mtb) infecting nearly a quarter of the world’s population ^1^. At the centre of TB pathology lies the granuloma, a dynamic immune structure that arises in response to infection. Composed of a core of differentiated macrophages surrounded by lymphocytes, the granuloma serves to restrict bacterial dissemination. Despite this protective role, granulomas often fail to completely eradicate the bacteria, creating a niche where Mtb can persist in a latent state. This dual nature—simultaneously containing the bacteria and enabling its survival—reflects the delicate balance between host defences and bacterial strategies. Understanding granuloma formation and function is therefore essential for elucidating TB progression, as it encapsulates the complex interplay between immune responses and pathogen persistence ^2^.

*In vitro* granuloma models have been instrumental in advancing our understanding of TB mechanisms and could be envisioned as a preclinical model to evaluate new anti-TB strategies ^3^. However, many of these models rely on inactivated commercial human AB serum (CHS) to supplement the cell culture medium ^4,5^, which may not fully replicate the complexity of the native immune environment. Autologous serum—either native autologous serum (NAS) or heat-inactivated autologous serum (HIAS)—has been suggested to better support certain immune cell functions, such as cytokine secretion, lymphocyte proliferation, and complement activity (Figure 1A) ^6,7^. These factors could potentially influence granuloma dynamics and immune polarization ^8,9^. Despite this possibility, the role of autologous serum in *in vitro* granuloma formation and its impact on immune response dynamics remains to be investigated.

**Figure 1:**
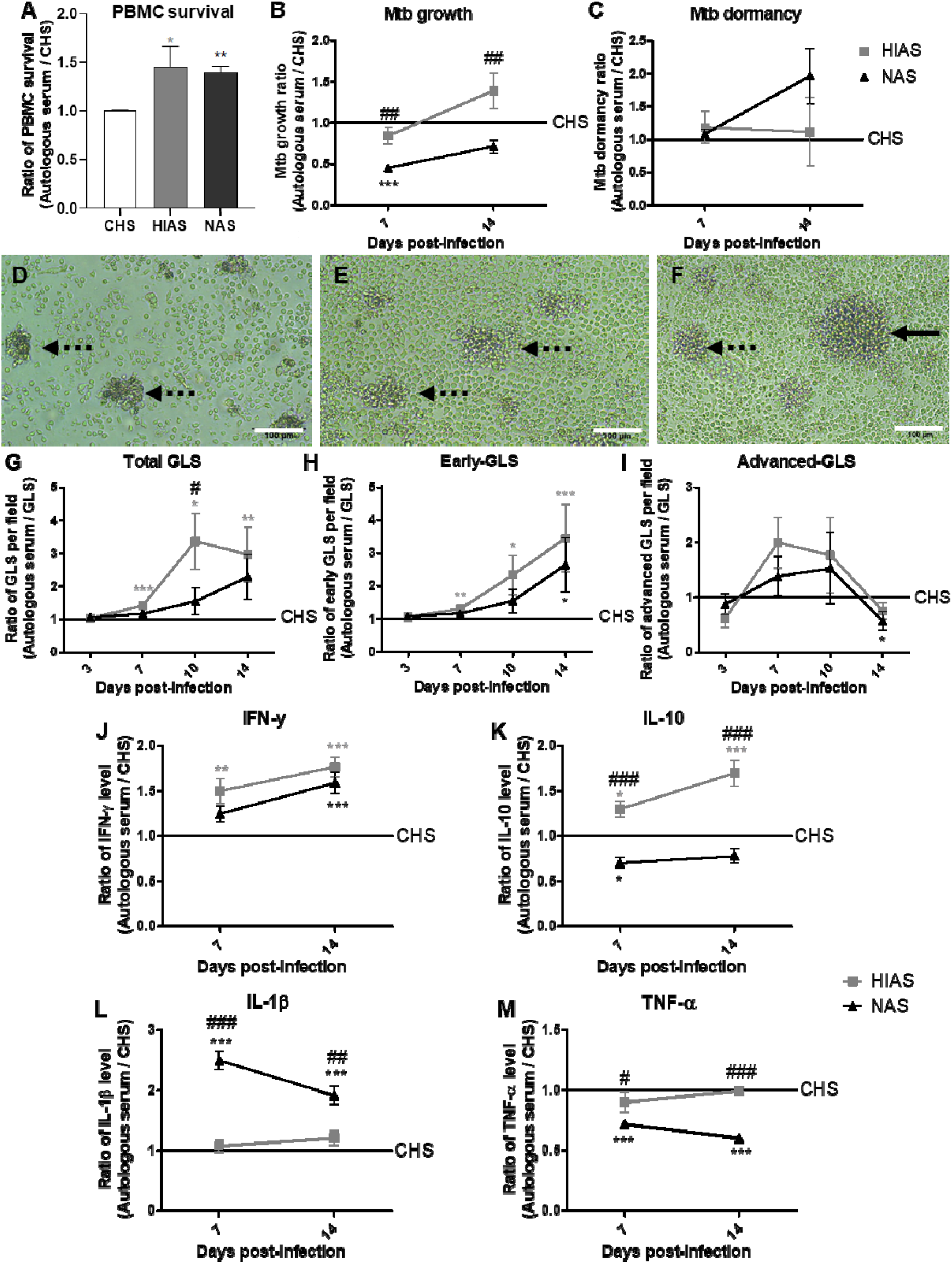
Impact of autologous serum on an *in vitro* granuloma model to study Mtb infection dynamics. **A**. PBMC survival after 7 days of culture in culture media supplemented with commercial human AB serum (CHS), heat-inactivated autologous serum (HIAS) or native autologous serum (NAS). **B-M**. PBMC were infected with the H37Ra Mtb reference strain at a multiplicity of infection of 1:10 (bacteria:cell), in presence of CHS (used as a control to normalise the results), HIAS (grey square) or NAS (black triangle), to induce spontaneous formation of *in vitro* granulomas. At 7- and 14-days post-infection, after granuloma disruption and cellular lysis, (**B**) Mtb growth was evaluated by CFU counting and (**C**) Mtb dormancy was assessed by quantifying differentially culturable bacteria in the presence or absence of resuscitation-promoting factors using MGIT (Mycobacterial Growth Incubator Tube) time to positivity system, reflecting the bacterial growth ^3^. (**D-F**) Representative bright field images of *in vitro* granulomas at 7-days post-infection from a representative donor. Dotte arrows show early grade GLS and complete one show advanced grade GLS. Scale bar = 100μm. (**G-I**) At 3-, 7-, 10- and 14-days post-infection, the dynamics of the total (**D**), early grade (**E**) and advanced grade (**F**) granuloma-like structure (GLS) observed per field was assessed by optical microscopy. Early grades correspond to start of granuloma formation, involving mainly monocytes/macrophages, with diameter <100μm. Advanced grades correspond to multilayer structures involving monocytes/macrophages surrounded by lymphocytes, with diameter >100μm ^10^. (**J**) IFN-γ, (**K**) IL-10, (**L**) IL-1β and (**M**) TNF-α release in cell culture supernatant was evaluated at 7- and 14-days post-infection, using Simple Plex Cytokine Screening Panel (Bio-techne, Minneapolis, USA) on a EllaTM (Bio-techne). Values for each condition are the mean ± standard deviation of four healthy donors and two independent experiments. Means were compared using One-Way ANOVA followed by Bonferroni correction. Grey stars represent the comparison between HIAS and CHS, black stars represent the comparison between NAS and CHS, # represent the comparison between HIAS and NAS. *p<0.05, **p<0.01, ***p<0.001, ****p<0.0001.

To address this, we assessed the dynamics of Mtb infection and granuloma formation using a previously developed protocol. Peripheral blood mononuclear cells (PBMC) from four healthy donors were infected with the avirulent Mtb strain H37Ra to induce spontaneous formation of *in vitro* granulomas, as previously shown ^3^. The experimental setup compared the effect of culture media supplementation with CHS, HIAS, and NAS on bacterial growth and dormancy, granuloma-like structures (GLS) formation, and immune response dynamics over 14-days post-infection (dpi). Bacterial growth was evaluated by CFU counting, bacterial dormancy was assessed by quantifying differentially culturable bacteria in presence or absence of resuscitation-promoting factors, GLS were quantified by optical microscopy, and immune response dynamics was analysed by measuring cytokine levels in culture supernatants ^3,10^. All results were normalized on those obtained for each donor upon infection in presence of CHS, as it is the widely used condition.

Our results showed that infection in the presence of different serum types had a modest impact on bacterial growth dynamics (Figure 1B). Growth was comparable upon infection in the presence of CHS and HIAS, while infection in the presence of NAS significantly reduced bacterial replication, with ratios of 0.45±0.042 and 0.71±0.077 compared to CHS at 7- and 14-dpi, respectively. Despite this, no significant difference was observed regarding induction of Mtb dormancy (Figure 1C). Regarding GLS formation (Figure 1D-I), infection in the presence of HIAS and NAS resulted in enhanced GLS formation compared to CHS, with up to 2.96±0.83 and 2.29±0.68 times more GLS per field observed at 14-dpi, respectively (Figure 1G). This was mainly due to the significantly higher formation of early-GLS (up to 3.44±1.02 time more early-GLS per field at 14-dpi, Figure 1H) and just a trend toward a higher formation of advanced-GLS at 7- and 10-dpi (Figure 1I). The marked increase in GLS numbers observed over time with autologous serum, whether heat-inactivated (HIAS) or native (NAS), is likely driven by improved PBMC survival in its presence. This hypothesis is supported by prior findings evaluating impact of culture media supplementation with autologous plasma ^11^ and our own observations (Figure 1A), indicating enhanced cell viability during culture with autologous serum, which may provide a more favourable environment for spontaneous and continuous GLS formation.

This higher formation of GLS upon infection in presence of autologous serum, whether HIAS or NAS, was associated with a higher production of IFN-γ compared to CHS (1.77±0.51 and 1.59±0.53 at 14-dpi, respectively, Figure 1J). However, the cytokine profiles diverged between these conditions. For infection in presence of HIAS, the parallel increase in IL-10 production suggested the preservation of a balanced pro- and anti-inflammatory response. In contrast, infection in the presence of NAS was characterized by reduced IL-10 production (down to a ratio of 0.70±0.27 at 7-dpi), indicative of an intensified pro-inflammatory response (Figure 1K). This heightened response was further evidenced by significantly elevated IL-1β levels (2.50±0.69 at 7 dpi and 1.92±0.68 at 14 dpi), despite a slight reduction in TNF-α production (Figure 1L-M). The active complement system in NAS likely contributes to this enhanced pro-inflammatory profile. The complement system is critical in modulating immune responses during Mtb infection, promoting inflammasome activation and IL-1β secretion ^12,13^. Additionally, the complement system enhances Mtb phagocytosis and intracellular killing by macrophages, and its dysfunction has been linked to increased susceptibility to TB in patients with type 2 diabetes mellitus ^13,14^. This mechanism could explain the reduced bacterial load observed upon infection in presence of NAS (Figure 1B). Overall, these distinct immune profiles underscore the differing impacts of the serum used on immune response dynamics.

In summary, our findings suggest that the type of serum used during infection has significant impact on both bacterial growth dynamics and GLS formation and the associated immune response. Notably, the use of native serum, containing the complement system with key-role in immune modulation and bacterial clearance, enables better control of the bacterial load, whilst the autologous serum favours GLS formation. This model offers potential as a preclinical tool for evaluating TB therapies, where physiological relevance is important, particularly with cells from TB patients. Incorporating the patient’s own serum allows the inclusion of host-specific components, such as cytokines, immune factors, and the complement system, which are essential for accurately reflecting the patient’s immune environment to mimic *in vivo* conditions, thereby advancing our ability to test and develop targeted therapies.

Despite these insights, further exploration is required to elucidate the precise mechanisms underlying these immune dynamics, including the roles of the complement system. Additionally, it is well-established that the H37Ra strain, used in this study, induces a type I immune response, unlike the virulent H37Rv strain, which favors a type II response ^15^. Characterizing this model with virulent Mtb strains will be critical for advancing its relevance as a preclinical tool.

In conclusion, autologous serum supports sustained GLS formation and offers a more representative environment for studying host-pathogen interactions compared to commercial human serum. While additional characterization is needed, our results underscore the importance of autologous serum for developing physiologically relevant *in vitro* granuloma models.

## SUPPLEMENTARY METHODS

### Ethics Declaration

Peripheral blood samples from 4 healthy donors were obtained from the Lyon Blood Bank (Etablissement Français du Sang, EFS, Lyon, France). In compliance with EFS procedures and Article R.1243–49 of the French Public Health Code, non-opposition to the use of blood for research purposes was obtained from all donors. Donor personal data were anonymized at the time of donation. Both sexes were included without discrimination, and the patients were considered healthy following an interferon-gamma release test after QuantiFERON-TB Gold test (Qiagen, Hilden, Germany), which was negative. Each experiment was conducted on four healthy donors in duplicate.

### Generation of *in vitro* granulomas

PBMCs were thawed and washed twice in RPMI with 10% fetal bovine serum. Trypan blue dye exclusion method was used to confirm sample viability greater than 95%. PBMC were adjusted at a density of 10^6^ cells/mL in RPMI supplemented with 10% of commercial human AB serum (CHS, Sigma-Aldrich), heat-inactivated autologous serum (HIAS) or native (not heat-inactivated) autologous serum (NAS) and seeded in 96-well culture plate coated with collagen at 10 μg/cm^2^ (Sigma-Aldrich) and fibronectin at 0.1 μg/cm^2^ (Sigma-Aldrich) for overnight recovery. PBMC were then infected with H37Ra bacterial suspension in log phase at a multiplicity of infection (MOI) of 1:10 (bacteria to cells) in RPMI supplemented with 10% of CHS, HIAS or NAS, at 37 °C in 5% CO_2_ atmosphere, to induce spontaneous formation of *in vitro* granulomas.

### Bacterial growth and dormancy assessment

At 7- and 14-dpi, all well contents were collected, and cells were recovered after coating digestion by a 15 min incubation at 37°C in the presence of collagenase 1 mg/mL (Thermo Fisher Scientific Inc, Rockford, IL, USA). Recovered cells were lysed with distilled water containing 0.1% Triton X100 (Sigma-Aldrich) for 10 min, followed by 3 min of vortex agitation in presence of 1-mm diameter glass beads to disrupt bacterial clumps.

To assess bacterial growth dynamics, mycobacteria were plated in duplicate on 7H10 agar supplemented with 10% OADC (Oleic acid, Albumin, Dextrose, Catalase; Becton Dickinson, Sparks, MD) and incubated for 3-4 weeks at 37°C. To assess bacterial dormancy, mycobacteria were inoculated in MGIT (Mycobacteria Growth Indicator Tube) for which a quarter of the medium volume was replaced or not by Mtb filtrate culture supernatant in stationary phase containing resuscitation promoting factors (RPF) ^16,17^. Analysis of the fluorescence was used to determine if the tube was instrument positive; i.e., the test sample contains viable organisms and results were expressed using MGIT time to positivity (TTP) system, reflecting the bacterial growth ^18,19^. Quantification of Mtb dormancy was assessed by analyzing the delta between TTP without and with RPF.

### Granuloma formation dynamics

Granuloma-like structure (GLS) formation dynamics was conducted by optical microscopy at 3-, 7-, 10-, and 14-dpi on 4 fields per condition from 4 healthy donors in duplicate. The granulomas were classified into 2 stages. Early stages correspond to start of granuloma formation, involving mainly monocytes/macrophages, with diameter <100μm. Advanced stages correspond to multilayer structures involving monocytes/macrophages surrounded by lymphocytes, with diameter >100μm ^10^.

### Cytokine release assay

At 7- and 14-dpi, cell supernatant was recovered and IFN-γ, IL-10, IL-1β and TNF-α release in cell culture supernatant was assessed using Simple Plex Cytokine Screening Panel (Bio-techne, Minneapolis, USA) on a EllaTM (Bio-techne).

#### Statistical analysis

Statistical analyses were performed with Graph Pad Prism 5. Data were normalized on those obtained with CHS, which is the most commonly used serum in *in vitro* granuloma models. Values for each condition are the mean ± standard deviation of four healthy donors and two independent experiments. Means were compared using One-Way ANOVA followed by Bonferroni correction. Grey stars represent the comparison between HIAS and CHS, black stars represent the comparison between NAS and CHS, # represent the comparison between HIAS and NAS. *p<0.05, **p<0.01, ***p<0.001, ****p<0.0001.

